# Leveraging protein dynamics to identify cancer mutational hotspots in 3D-structures

**DOI:** 10.1101/508788

**Authors:** Sushant Kumar, Declan Clarke, Mark B. Gerstein

## Abstract

Large-scale exome sequencing of tumors has enabled the identification of cancer drivers using recurrence and clustering-based approaches. Some of these methods also employ three-dimensional protein structures to identify mutational hotspots in cancer-associated genes. In determining such mutational clusters in structures, existing approaches overlook protein dynamics, despite the essential role of dynamics in protein functionality. In this work, we present a framework to identify driver genes using a dynamics-based search of mutational hotspot communities. After partitioning 3D structures into distinct communities of residues using anisotropic network models, we map variants onto the partitioned structures. We then search for signals of positive selection among these residue communities to identify putative drivers. We applied our method using the TCGA pan-cancer atlas missense mutation catalog. Overall, our analyses predict one or more mutational hotspots within the resolved structures of 434 genes. Ontological and pathway enrichment analyses implicate genes with predicted hotspots to be enriched in biological processes associated with tumor progression. Additionally, a comparison between our approach and existing hotspot detection methods that use structural data suggests that the inclusion of dynamics significantly increases the sensitivity of driver detection.

## Introduction

Large-scale cancer genome studies such as The Cancer Genome Atlas (TCGA) project^1,2^ and the International Cancer Genome Consortium (ICGC)^3,4^ have generated comprehensive catalogs of somatic alterations for various cancer cohorts. The majority of these somatic variants incur little or no functional consequence on tumor progression, and are thus often termed neutral ‘passengers.’ In contrast, a handful of ‘driver’ mutations are considered to provide a selective advantage to cancer cells. One of the critical goals of TCGA and ICGC projects has been to distinguish between these positively selected “driver mutations”^5–7^ from a large number of neutral passenger mutations.

A majority of the cancer driver detection algorithms quantify the recurrence of mutations to identify significantly mutated genes and non-coding genomic elements^8–11^. However, the somatic mutation landscapes of cancer genomes are highly heterogeneous^12–14^ and exhibit a long tail of low-frequency mutations^11,13,15–17^. The presence of this long tail of rare somatic mutations, along with limited cohort sizes, makes recurrence-based driver identification very challenging. An alternative is to employ algorithms that aggregate mutation recurrence on gene/element-levels^18,19^ or to predict the molecular functional impact of mutations^20^ to distinguish drivers from passengers. Compared to protein-truncating mutations and large structural variants, missense mutations induce subtle changes, which are often difficult to interpret on the phenotypic level. Thus, identifying missense driver mutations based on their molecular functional impact is also challenging. In contrast, the signal of positive selection aggregated on functional elements or sub-regions of the coding genome (such as protein domains^21–23^, post-translational modification sites (PTMS)^24–26^, protein interaction interfaces^27,28^ and mutation cluster/hotspots^29–31^) has been shown to be effective, despite their intrinsic limitations.

Prior studies have identified driver mutations based on their presence in mutational clusters^29–31^, which are sometimes called “hotspot” regions. These mutational clusters are defined based on the proximity of somatic mutations within the primary sequence^29,31^ or three-dimensional structure of a given protein^32–36^. Sequence-based mutation cluster identification algorithms^29,31,37^ discover significantly mutated genes while considering an appropriate background mutation model, trinucleotide context of mutations and distribution of silent mutations. However, sequence-based approaches miss many hotspot regions, as they ignore spatial proximity between residues that may be far apart in sequence but can be very close in 3-dimensional(3D) space^38,39^, in the context of the fully-folded protein or protein ensembles. In contrast, despite being inherently limited due to incomplete structural coverage of the proteome, 3D structure-based mutational cluster definitions provide physical intuition or mechanistic insight into the roles of a mutational cluster in cancer progression^32–36^. These structure-based methods compute residue distances or generate residue-residue contact networks in the 3D structures of proteins to identify a group of spatially proximal residues. Furthermore, mutation shuffling is performed to identify significantly mutated residue clusters or hotspots on a protein structure. However, it is important to note that, current approaches under this framework have failed to consider protein dynamics.

Proteins are inherently dynamic bio-molecules and sample large ensembles of conformations^40–43^. The energy landscape underlying the distribution of structures in these ensembles are often altered based on external (thermodynamic)^44,45^ or internal (allosteric) signals^43,46^. Previous biophysical studies have clearly shown the crucial role of protein motions in conferring protein functionality^47^. Thus, one could argue that prior structure-based driver detection methods that employ only the static structure of proteins are less sensitive when attempting to identify functional residues through the mutation clustering approach.

In particular, a static crystalized structure provides only one limited snapshot of the protein, most likely close to (or at) the bottom of the free energy landscape. In contrast, motion-weighted community detection approach better reflects physical reality where proteins undergo two general types of dynamics. First, a protein can dynamically oscillate around the bottom of the energetic well or in second type of dynamics the underlying free energy landscape changes in distinct ways, thereby shifting the protein conformation to an alternative functional state. In each of these scenarios, communication between different communities plays a pivotal role in the proper functioning of the protein. We posit that hotspot communities exist in large part because certain select communities either play especially essential roles in these functional dynamics or because their contributions to such dynamics are especially sensitive to mutations. Static representations of protein structures presumably fail to define communities in light of their essential roles in dynamics, and thus function. Furthermore, they potentially miss many critical mutational clusters with a potential role in cancer progression.

In the current work, we address this issue by explicitly incorporating protein dynamics into our new framework to identify mutational hotspot communities in protein structures. We applied this framework to the TCGA pan-cancer atlas catalog of missense mutations to identify genes with significantly mutated residue communities in protein structure. Our pan-cancer analysis identifies 424 unique genes with at least one hotspot community in the corresponding protein structure. The majority of these genes are involved in critical biological processes and pathways involved in cancer progression including DNA repair, signal transduction, immune response, apoptosis, and post-translational modifications. As expected, we observe higher cross-species conservation score and greater functional impact scores for mutations present in these hotspot communities. Furthermore, our prediction includes previously characterized driver genes with hotspot communities in corresponding protein structure. Additionally, we also identify novel genes with at least one hotspot community that were not detected by other mutation cluster algorithms lacking protein dynamics information. Finally, we highlight some examples of driver genes containing hotspot communities which are predicted to play a vital role in cancer progression.

## Material and Methods

### SNV dataset and mapping onto protein structure

In this study, we leveraged the MC3(multiple-center mutation calling in multiple cancer)^48^ somatic mutation dataset generated as part of the TCGA pancan atlas project. Briefly, the MC3 call set was generated using approximately 10,000 tumor/normal whole exome sequences belonging to 33 different cancer types. Multiple callers, including MuTect^49^, RADIA^50^, SomaticSniper^51^, and VarScan^52^ were applied to obtain high-confidence variant calls. Subsequent filtering removed mutations due to lack of coverage, potential germline contamination, and other artifacts. We utilized version 2.8 of the publicly accessible MC3 variant call set^5^. Furthermore, we only analyzed missense mutations that were designated as ‘PASS’ based on the filtering criterion. Moreover, we only analyzed variants from samples that were included in the whitelist samples and were not hyper-mutated. This subset comprises 2.85 million mutations from 8937 samples in the pancan atlas project. Approximately 2.29 million mutations in this subset occupy the coding regions of the genome that consists of 1.5 million missense mutations, 1.18 million silent mutations, 0.6 million nonsense mutations, and 3.7K splice mutations.

We applied the Variant Annotation Tool (VAT)^53^ to map TCGA missense mutations onto protein structures. For each missense mutation, VAT provides an annotation that includes gene name, transcript name, and the position of the residue getting affected in the translated protein sequence. Additionally, it also provides the residue identity of the original and mutated residues. Subsequently, we integrated VAT annotations with a BioMart^54^ derived identifier map, which consists of the gene identifier, transcript identifier, and the corresponding PDB ID, if available. We restrict our analyses to mutations that map to crystal structures with resolution better than 3.0 Å. This restriction was applied to in order to most precisely identify residue communities in protein structures. Overall, we mapped 0.329 million missense mutations on approximately 17,300 crystal structures in the current study.

### Workflow to identify three-dimensional hotspot communities in cancer

As discussed above, our framework to predict driver genes through identification of hotspot communities is novel compared to prior approaches as we explicitly include protein dynamics information in our workflow (**Fig 1)**. Briefly, our integrative workflow includes three distinct components. First, we model large-scale conformational changes of a protein to identify dynamic sub-regions of proteins (or “communities”). The large-scale conformational changes are modeled using anisotropic network models (ANMs)^46,55^. Subsequently, we model protein structure as a residue-interaction network, where each residue constitutes a node in the network, and edges (or connections between these nodes) form the physical interactions between these nodes. Furthermore, edges in a network can be ‘weighted’ using the extent to which contacting residues exhibit correlated movements within the dynamic structure of the protein. Highly correlated motion (or movement vectors) between two residues that are physically in contact (though not necessarily covalently linked) suggest that knowledge of the motions for one residue can provide a great deal of information regarding the motions of the other residue. This mutual knowledge, in a sense, suggests a strong degree of informational flow between residues. The weight for each edge in the network corresponds to the “effective distance” of this edge, in which a strong degree of correlated motion results in a short distance, and a weak correlation in the motions results in a long distance. With this motion-weighted protein network, communities of resides are defined with the Girvan-Newman algorithm^56^. Communities are then defined as residue groups in which each residue of a given community is connected to other residues of the community, and only tangentially connected to residues outside the immediate community. These network-weighted communities thus form densely inter-connected neighborhoods.

**Fig 1.**
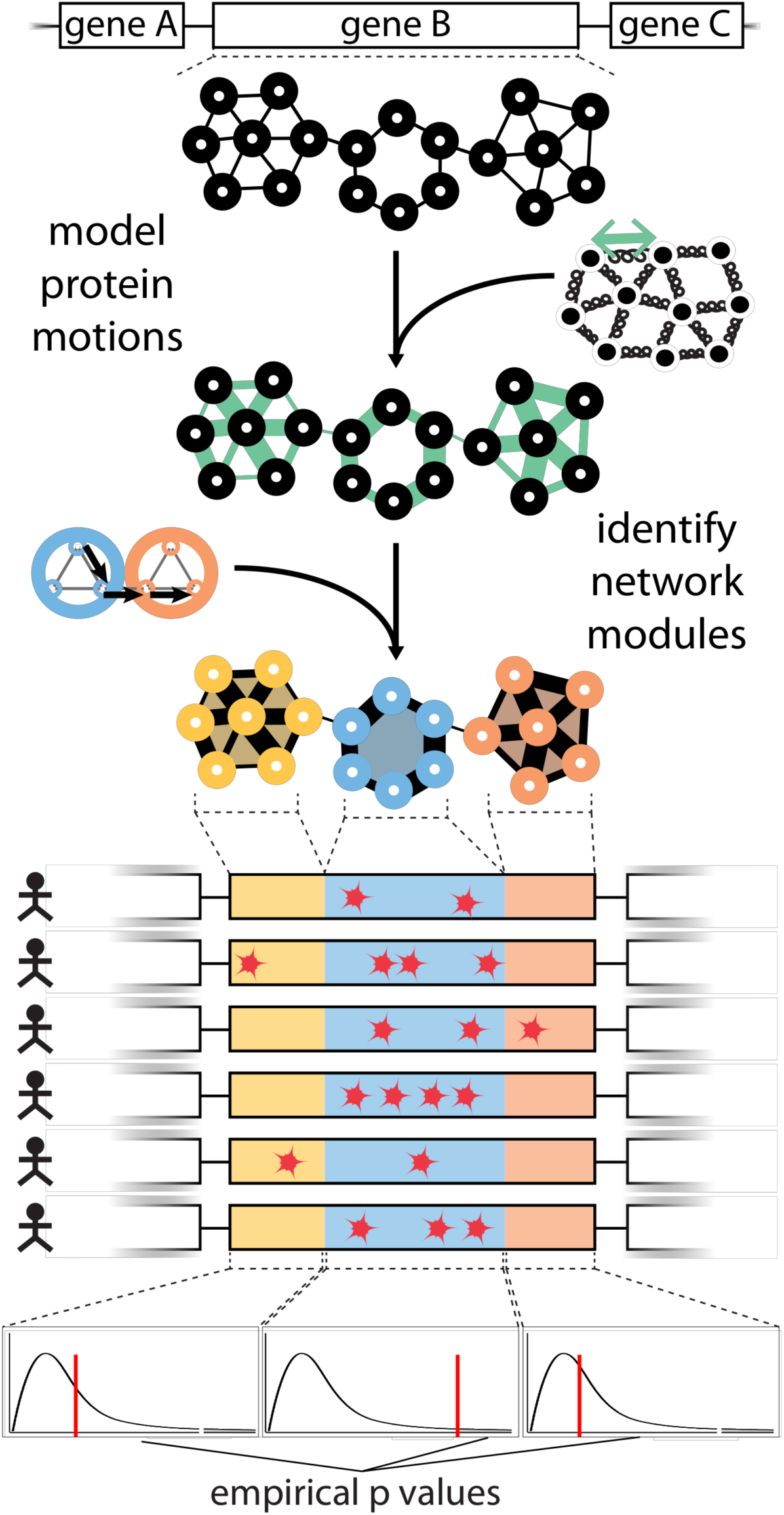
Workflow of HotCommics to identify putative driver genes: This integrative approach utilizes protein community information along with mapped mutations onto protein structure to identify significantly mutated communities in protein structure. Fisher method is employed to quantify significance value for each community with mapped mutations.

In order to identify mutational hotspot communities on a given protein structure, we mapped missense mutations from TCGA cohorts onto three-dimensional protein structures. Subsequently, we computed the frequency of mapped mutations for each community on the pan-cancer level as well as in specific cancer cohorts. Furthermore, for each community with mapped mutations, we performed a Fisher exact test to determine whether variants fall within a given community is more frequently mutated than what would be expected by chance. This significance test assigns an empirical p-value, which we correct for multiple hypothesis testing using the Benjamini Hochberg method to identify significantly mutated hotspot communities on protein structure for a given gene. We note that, for a substantial number of genes, there are multiple PDB structures available. We remove this structural redundancy using structural coverage (highest fraction of residues covered in the structure) as a filter to provide one to one mapping between PDB structure and corresponding gene. The source code for the workflow is available on the project’s Github page (https://github.com/gersteinlab/HotComms).

### Downstream Analyses

We performed many downstream analyses to further validate our predictions. We extracted PhyloP^57^ and CADD^58^ score for each mutation mapping onto protein structures. Furthermore, we classified mutations into hotspot and non-hotspot mutations based on whether mutations are mapped onto residues belonging to hotspot communities or otherwise. Subsequently, we compared the phyloP score and CADD score distributions for hotspot and non-hotspot mutations. We performed two-sided Kolmogorov-Smirnov(KS) test to assess the significance of conservation score differences between hotspot and non-hotspot mutations. We apply the same method to quantify such disparities for the molecular functional impact (CADD) score for hotspot and non-hotspot mutations. Here, our null hypothesis is that the conservation or impact score for hotspot and non-hotspot mutations are on average not different as they are being drawn from the same distribution.

We also performed gene ontology(GO) enrichment and pathway enrichment analyses to further validate the role of our putative driver genes in tumor progression. For the GO analysis, we calculated the enrichment based on biological processes available from the GO database^59^, and we performed pathway enrichment analysis using the Reactome^60^ as well as the KEGG database^61^. We visualized the enrichment analysis result using the clusterProfiler^62^ package available in Bioconductor.

Additionally, we also compared our predicted driver gene list derived from our hotspot community analysis with other approaches that detect driver genes based on the presence of mutation clusters on sequence or structure levels. One of the key differences between our approach and other approaches is that we employ information on protein dynamics (along with structural data) to determine hotspot communities. For structure-based methods, we obtained driver gene list predicted from HotSpot3D^35^, 3DHotSpot^34^, HotMap^36^ algorithms. All three of these algorithms were previously applied on the TCGA Pancan Atlas data^5^, which allows us to make meaningful comparisons with our work. However, we also note small differences in our workflow compared to other structure-based approaches. For instance, HotMap tools employ homology-model derived structures compared to other methods that rely only of experimentally determined structure. Moreover, our method was applied only on crystal structure at higher resolution compared to other methods that included NMR as well as crystal structures at higher resolution. Finally, we also employed predicted driver genes from sequence-based cluster analysis tool (OncodriverClust^31^) and previously curated driver genes in the cancer gene census(CGC) database^63,64^. We note that we excluded driver genes in CGC that play role in cancer through INDELs, copy number aberrations or other structural variations. We used UpsetR^65^ package in R to visualized the multiway comparisons among predicted driver genes from various tools and CGC database.

Finally, we also performed gene expression analysis to validate the role of our putative driver genes in cancer at the transcriptome level. For this analysis, we obtained the TCGA RNA-Seq quantification available for samples in the Pancan Atlas project^2^. For each gene in our putative driver gene list (based on hotspot community information), we compared the gene expression distribution for sampled that harbored missense mutations to those that are not mutated. We performed a two-sided KS test to evaluate the significance value for each gene in our putative gene list. These significance tests were carried out separately for each cancer-type. However, we combined the significance level(p-value) for each gene across multiple cancer types using the Fisher method. We visualized significantly differentially expressed genes using a standard QQ plot.

## Results

### Pan-cancer analysis of genes containing mutations clusters

We applied our workflow to identify significantly mutated hotspot communities for each cancer cohort as well as on the pan-cancer level. As expected, we observed a comparatively higher number of genes with at least one hotspot community on the pan-cancer level compared to cancer-specific analysis. Our pan-cancer analysis identifies hotspot communities present on protein structures of 434 unique genes (**Fig 2a, supplement table S1**). In contrast, a cancer-specific analysis revealed 56 potential driver genes with 186 significantly mutated hotspot community in the corresponding protein structure (**Supplement table S2**). Some of these genes (including TP53, PIK3CA, BRAF, SPOP, KRAS, HRAS, and PTEN) have been previously shown to be a driver for different cancer types. However, we also identified numerous novel genes containing hotspot communities that might drive cancer progression. Previous studies suggest that some of these novel genes including RHOC, NCOA1, and KLHL12 are involved in various signaling pathways. Similarly, PSPC1, FOXO3, and XRCC5 are known to be pivotal for immune response, apoptosis, and DNA repair, respectively. Furthermore, among these 434 genes, 12 genes had five or more hotspot communities whereas 352 genes had just one hotspot community on their corresponding protein structure. These observations highlight the efficacy of our approach in identifying novel and low-frequency putative driver genes with hotspot communities.

**Fig 2.**
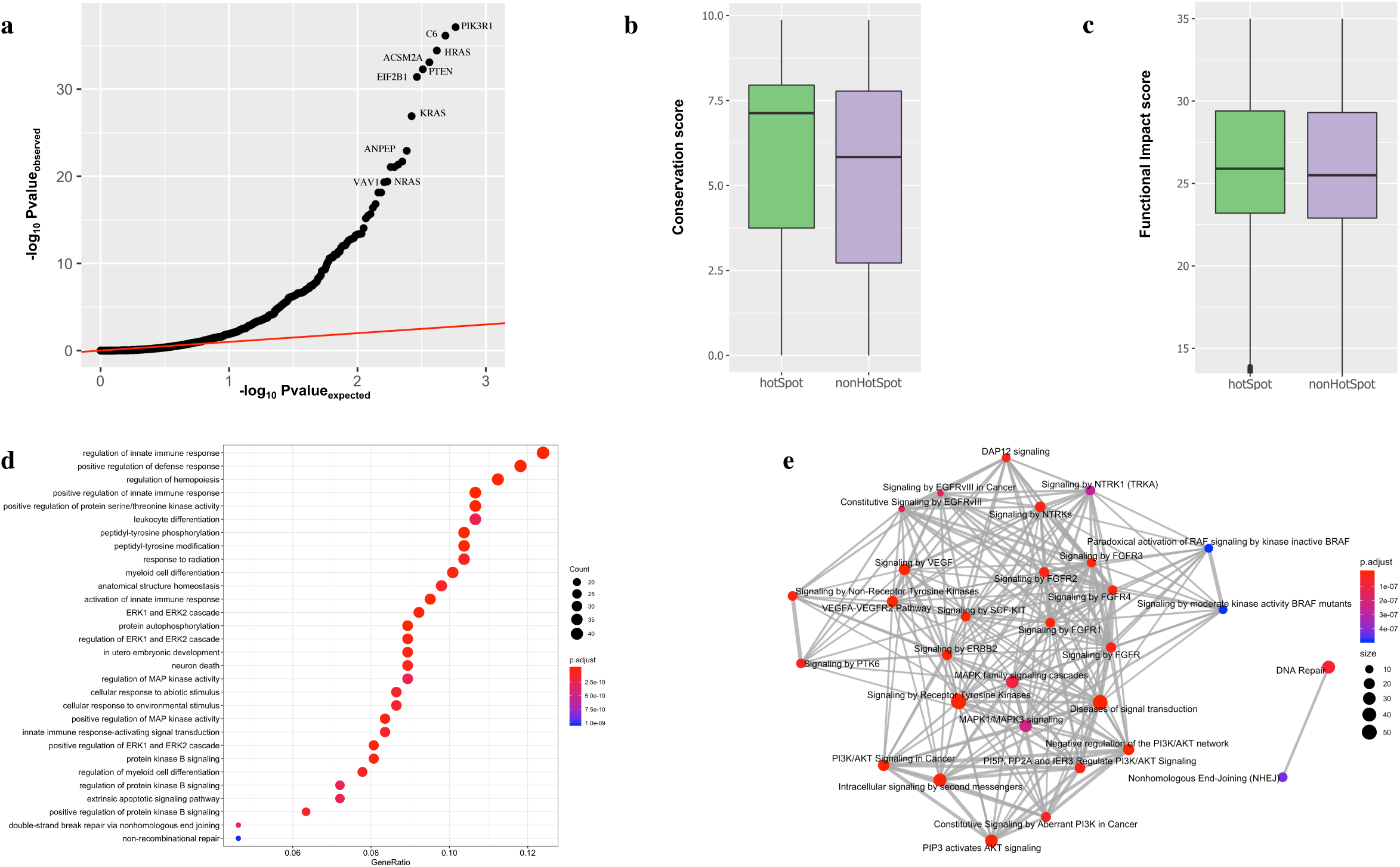
Pan-cancer analysis of putative driver genes with hotspot communities: a) pan-cancer q-q plot for genes with hotspot communities, b) PhyloP conservation score comparison between mutations occupying hotspot communities against non-hotspot communities on protein structures, c) CADD score correlation between mutations occupying hotspot communities against non-hotspot communities on protein structures, d) Biological process enrichment analysis for putative driver genes with at least one hotspot. X-axis corresponds to gene ratio that corresponds to the fraction of putative driver genes belonging to a particular biological process. The color code and size correspond to corrected p-value and number of genes involved in the biological process, respectively, e) Reactome based pathway enrichment analysis. The color code and size correspond to corrected p-value and number of genes involved in the biological process, respectively.

Mutation cluster-based approaches assume that residues constituting such clusters are essential for protein functions. Thus, a majority of cancer missense mutations occupying these hotspot communities are very likely to disrupt the protein function. In order to validate this assumption, we quantified the cross-species conservation measure (PhyloP score^57^) for mutations in hotspot as well as non-hotspot communities on protein structures. As expected, we observe higher average conservation score for mutations mapping to residues in hotspot communities compared to those, which are present outside. Furthermore, the observed difference in conservation is statically significant (two-sided KS test, p-value < 2e-5) (**Fig 2b)**. Similarly, the putative molecular functional impact (CADD score^58^) of mutations occupying hotspot communities was significantly higher compared to those mapping to non-hotspot communities (two-sided KS test, p-value < 2e-5) (**Fig 2c)**.

We also preformed gene ontology^62^ and pathway enrichment analysis to decipher the biological function of genes with predicted hotspot communities. The biological process based gene ontology enrichment analysis indicate role of putative driver genes in diverse biological function including immune response, cell differentiation, kinase activities, post-translational modifications, apoptosis and DNA repair (**Fig 2d & Supplement table S3)**. Similarly, reactome pathway^60^ based enrichment analysis suggest role of putative driver genes with hotspot communities in various signaling pathways (**Supplement table S4**) including NTRK signaling, DAP12 signaling, EGFR signaling and MAP kinase-associated signaling. Additionally, these genes are also enriched among DNA repair and non-homologous end-joining associated pathways (**Fig 2e)**. Furthermore, KEGG pathway^66^ based enrichment analysis indicate role of our putative driver genes in various cancer subtypes (bladder, pancreatic, breast, CML, melanoma, AML, glioma) (**Supplement Fig1 & Supplement table S5)**.

### Comparison of 3D structure based clustering methods

We performed consensus analysis between our approach to the driver genes curated in the COSMIC^67^ database. Furthermore, we also performed a comparison between putative driver genes identified using our workflow and genes identified as drivers by other mutation cluster detection algorithms that do not take protein dynamics into account. The majority of these additional algorithms employ the three-dimensional structure of a protein to identify mutational cluster except the OncoDriveClust^31^ tool, which searches for hotspot mutations on the sequence level. Overall, our workflow identified many additional genes (288 genes) with hotspot communities compared to other mutation hotspot analysis tools (**Fig 3a**). One exception being the HOTMAP^36^ algorithm that utilizes protein homology model in addition to protein structure. Thus, it identifies significantly higher number of unique genes (620 genes) with mutation cluster compared to any other tool. Furthermore, our approach identified 146 genes (34% of our gene list) with hotspot communities that are either curated as a driver gene in COSMIC or predicted to contain a mutation cluster by another tool (**Fig 3a**). Among these 146 genes, 89 genes overlapped with putative driver genes identified by HOTMAP algorithm, whereas 63 genes overlapped with drivers in COSMIC. As expected, we observed the lowest overlap (33 genes, 7% of our putative driver gene list) with sequence-based method (OncoDriveClust; **Fig 3a**).

**Fig 3.**
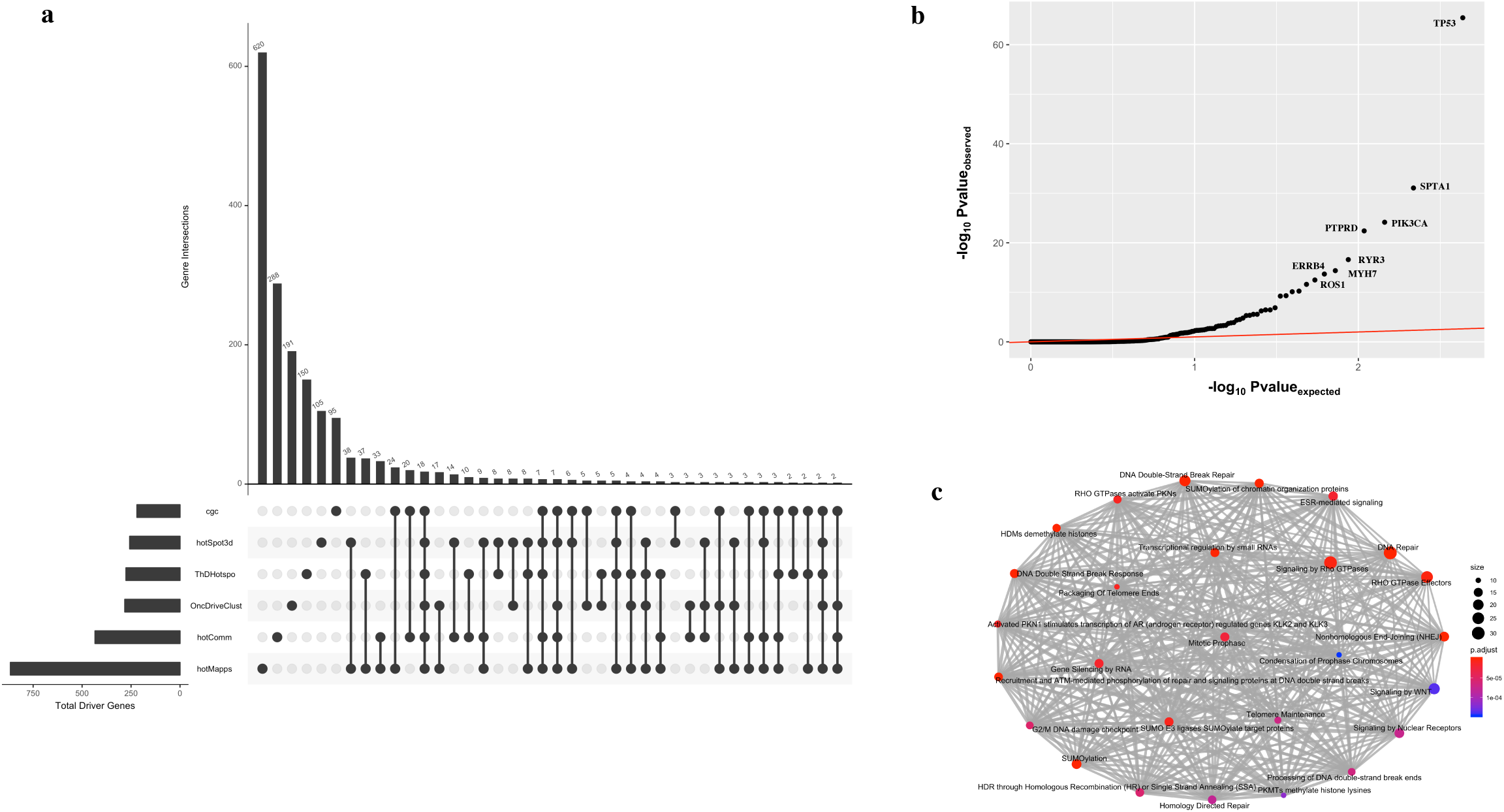
Pan-cancer analysis of putative driver genes with hotspot communities: a) Comparison of multiple driver detection algorithms including HotCommics. We used the most recent version of the Cancer Gene Census database for this analysis. Remaining algorithms were also run on the MC3 variant call set, b) Q-q plot highlighting differentially expressed putative driver genes across multiple cancer types, c) Pathway level enrichment analysis of singleton genes identified by the HotCommics algorithm that was novel for putative driver genes identified by other algorithms and CGC database.

Additionally, we analyzed TCGA expression data to obtain additional evidence corroborating the biological validity of putative driver genes identified through our workflow. Intuitively, one would expect a significant difference in gene expression level between samples with and without mutation for genes that were predicted to contain a significantly mutated hotspot community. For each candidate gene, we quantified the statistical significance in expression distribution differences using two-sided KS test. Furthermore, we performed this test for individual cancer type, and the corresponding p-values were combined across cancer types using Fisher’s method to provide a pan-cancer significance measure. Overall, our analysis identified 60 genes including TP53(p-value 3.59e-66), SPTA1 (p-value 8.58e-32), PIK3CA (p-value 7.06e-25), KRAS (p-value 5.73e-11), and EGFR (p-value 2.78e-06) that were differentially expressed across cancer types (**Fig 3b & Supplement table S6**). A subset of these differentially expressed genes such as MYH7 (p-value 4.22e-15), ROS1 (p-value 3.26e-13), TIAM1 (p-value 2.48e-12), PTPRD (p-value 3.96e-23), and HUWE1 (p-value 4.84e-10) are potentially novel driver genes with predicted hotspot communities (**Fig 3b & Supplement table S6**). Moreover, we note that 76% of our putative driver gene list with significantly mutated hotspot communities were differentially expressed in at least one TCGA cancer cohort.

Finally, we performed GO and pathway enrichment analysis on novel genes that we predict to contain mutational hotspot communities. However, these genes were neither present in the COSMIC driver database nor were predicted to encompass mutation cluster through other hotspot identification tools. We observed significant enrichment of these genes in crucial biological processes (**Supplement table S7**) including DNA conformation change, regulation of immune response, regulation of stem cell differentiation, nucleosome organization, and endothelial cell apoptotic process (**Supplement Fig2**). Similarly, pathway enrichment analysis implicates their role in DNA repair, SUMOylation, RHO GTPase activity, telomere maintenance, and various signaling pathways (**Fig 3c & Supplement table S8**).

### Case studies highlighting the roles of hotspot communities in deciphering driver mechanisms

Integration of protein 3D-structure and protein dynamics to identify driver genes has a clear advantage over other methods that do not leverage protein structure or protein dynamics information. Our method allows us to investigate disruption in protein structure and function induced by missense mutations that occupy within predicted hotspot communities. We also note that the majority of our hotspot communities encompass residues that are pivotal for important protein functions including allostery, bimolecular signaling, protein binding, and post-translation modifications. The sensitive detection of functional sites on protein structure helps to decipher the underlying biophysical mechanism that plays a crucial role in cancer growth. Here, we highlight three examples testifying the utility of our framework in gaining biophysical insight into cancer progression through disruption of predicted hotspot communities. These examples include an oncogene(BRAF), tumor suppressor gene(PIK3R1), and a novel putative driver gene(PTPRD) that are predicted to contain multiple hotspot communities on their respective protein structure.

### Missense hot spot communities: PIK3R1

The PI3KR1 gene encodes the alpha subunit of the enzyme Phosphatidylinositol 3-kinase regulatory, which plays a crucial role in a variety of cellular processes including cell survival, regulation of gene expression, cell metabolism and cytoskeletal rearrangement^68^. Mutations in PIK3KR1 gene has previously been implicated as a tumor suppressor gene in breast cancer. Recent therapeutic studies have targeted PI3K inhibition resulting in a decrease in cellular proliferation and reduced metastasis in the mouse model. PI3Ks are obligate heterodimers composed of a p110 subunit and a regulatory subunit. Previous studies have identified four distinct domains belonging to the catalytic P110 alpha subunit that harbor somatic mutations leading to an increase in PI3K activity. We observe two distinct hotspot communities (**Fig 4a)** on the co-crystal structure (PDB ID: 2V1Y) of the protein complex that compromises ABD domain of the P110 alpha subunit and the iSH2 domain of the p85 alpha regulatory subunit. The two hotspot communities are composed of 28(community 5) and 26(community 7) residues, respectively (Fig 4a). On the pan-cancer level, we observe 24 and 16 mutations that map to community 5 and community7 on the co-crystal structure, respectively. These distinct hotspot communities are adjacent to each other in the same helical structure. However, we observe a small kink in this helical structure, which presumably lead to distinct protein motions associated with these two different hotspot communities.

**Fig 4.**
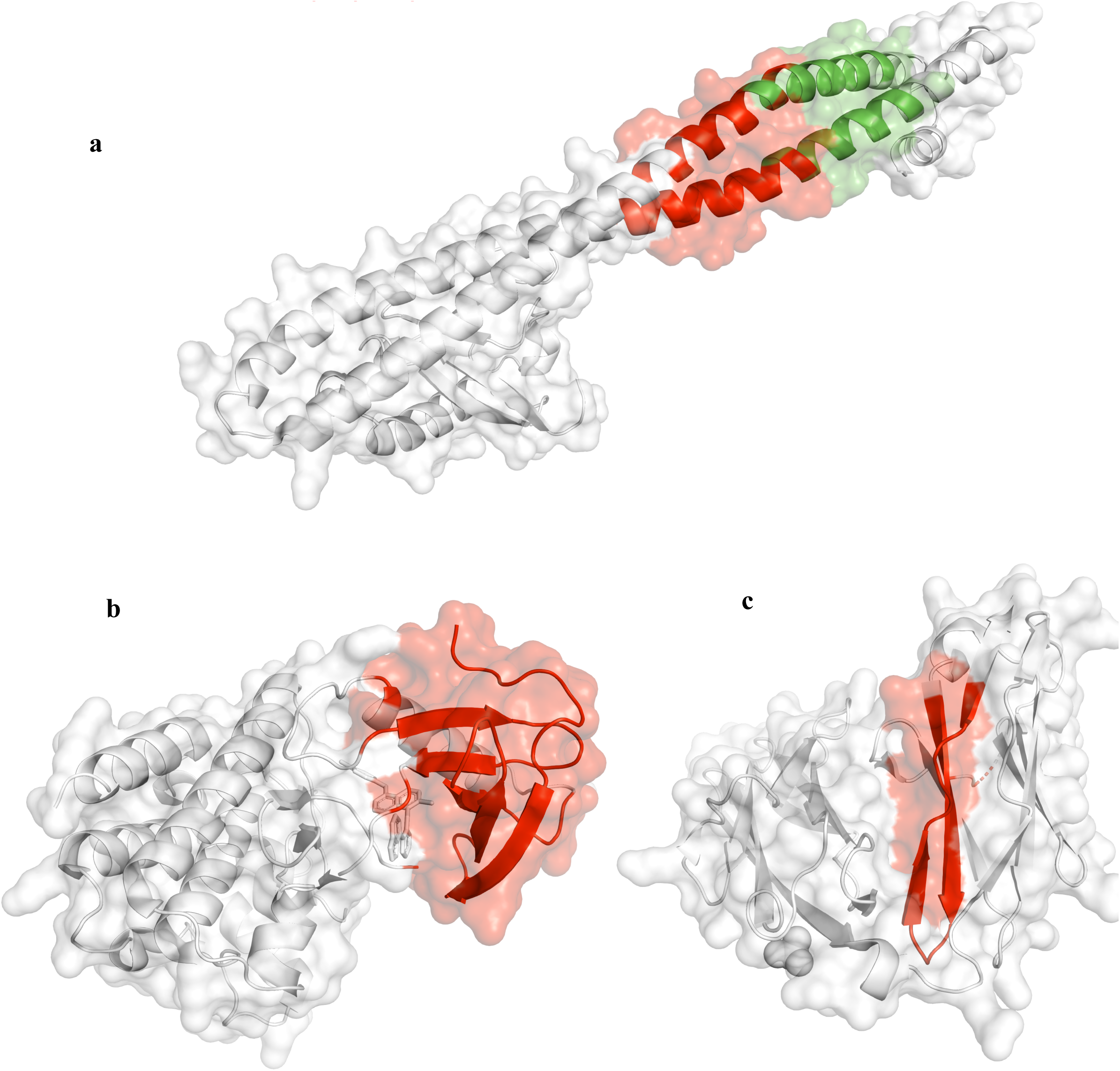
Examples of TSG, oncogene, and putative driver genes with hotspot communities: a) Example hotspot communities (shown in red) on the PIK3R1 gene as identified by our workflow. We note that previous studies have identified the PIK3R1 gene as a tumor suppressor gene, b) Example hotspot communities (shown in red) on the BRAF gene as identified by our workflow. We note that previous studies have identified BRAF1 gene as an oncogene, c) Example hotspot communities (shown in red) on the PTPRD gene as identified by our workflow. We note PTPRD is an example of novel putative driver genes with hotspot community with significant differential gene expression.

### Missense hotspot communities in BRAF gene

BRAF gene encodes a protein belonging to the serine/threonine protein kinase family that regulates MAP kinase and ERK signaling pathway^69^. This pathway is considered to be essential for a number of biological functions including cell differentiation, cellular growth, senescence, and apoptosis. Somatic mutations in the BRAF gene are often implicated in various cancer subtypes including melanoma, colorectal cancer, prostate cancer, non-small-cell lung cancer, and papillary thyroid tumors. It has been proposed that BRAF induce dysregulation in the binding of Ras proteins to Raf and MEK proteins in the Ras/RAF/MEK/ERK signaling cascade that leads to over-activation of the signaling pathway and subsequent oncogenesis. Multiple enzyme inhibitors have been designed to target BRAF kinase in the tumor. One such inhibitor SB-590885 has been co-crystallized with BRafV600E kinase domain at the X-ray resolution of 2.9 Angstrom (PDBID: 2FB8)^70^. A previous study indicates the role of pi-stacking interactions, hydrogen bonds and salt bridges in stabilizing the interaction between these two subunits in the crystal structure. In our study, we identified one hotspot community in this co-crystal structure (**Fig 4b**). This hotspot community is composed of 52 residues that constitute a beta sheet secondary structure. Interestingly, we also observe that SB-590885 inhibitor occupies the same hotspot community.

### Missense hotspot community in TPRD gene

The PTPRD gene encodes a protein that belongs to the protein tyrosine phosphatase(PTP) family. PTP proteins are considered essential for regulating cellular proliferation, differentiation, and oncogenic transformation. PTPRD gene encodes a transmembrane protein containing a cytoplasmic tyrosine phosphatase domain. Previous studies have shown that PTPRD genes are frequently deleted in various cancer types including glioma, neuroblastoma, and lung cancer^71^. However, we note PTPRD is not identified as missense driver in cosmic catalog. Moreover, previous studies did not identify presence of mutational hotspot communities in the PTPRD gene. In contrast, our analysis identifies one hotspot community in the crystal structure (PDB ID: 2YD7) of the receptor protein tyrosine phosphatase(RPTP) sigma subunit. RPTPs are cell surface proteins with intracellular PTP activity and extracellular domains that are sequentially homologous to cell adhesion molecules. Moreover, RPTP sigma subunit is considered necessary for nervous system development and function. In our analysis, somatic mutations mapped to two communities (community 2 & 4) on the crystal structure of the RPTP sigma subunit. Our workflow predicts one hotspot community that comprise of 47 residues in the crystal structure of PTPRD (**Fig 4c)** and adopts a beta strand conformation.

## Discussion

The underlying heterogeneous characteristic^72^ of cancer makes interpretability of genomic alterations in a cancer genome very challenging. In particular, genomic heterogeneity poses a major challenge in identifying key driver mutations in cancer. Large-scale cancer genome sequencing efforts have helped us to generate comprehensive catalogs of driver mutations^5^ in various cancer types. However, the canonical recurrence-based driver detection algorithms have failed to identify low-frequency or rare drivers. The limited cohort size^11^ and heterogeneity^14^ in cancer genome provides limited power to identify low-frequency drivers using the canonical position level recurrence algorithms. A simplistic approach to address the issue of missing rare driver will be to sequence more patients for a given cancer type. However, this approach will be particularly challenging for highly heterogeneous cancer cohorts with multiple subtypes^73^ within a cancer type. Moreover, this approach will not be practical for certain rare cancers including neuroblastoma, angiosarcoma, Hodgkin’s lymphoma, and others. A suitable alternative is to quantify recurrence over functional elements or sub-gene levels^74^ such as post-translational modification sites (PTMS)^25,26^, protein interaction interfaces^28^ and mutational clusters^33–36,38^. In particular, many driver detection algorithms search for the presence of mutational hotspot on the 3D-protein structures to identify putative driver genes. Compared to sequence-based driver detection methods, using protein structural data can help to decipher the underlying molecular mechanisms that influence cancer progression. However, current approaches to identify cancer mutation hotspots on protein structure and corresponding driver genes completely ignore the role of protein dynamics, which is considered essential for protein function. Thus, here we propose a new framework that utilizes protein dynamics along with the 3D-structure of proteins to identify missense hotspot communities on protein structure and corresponding putative driver genes.

Overall, our workflow identified 802 hotspot communities on crystal structures of proteins corresponding to 434 unique genes on the pan-cancer level. We also compared our putative driver gene list with previous experimental and prediction studies derived driver gene list. Among our putative driver gene list, we find 36% of genes are either known or predicted to be driver genes based on previous studies. We term the remaining 64% of genes as novel drivers in our study. We performed many downstream analyses on our putative driver genes to highlight their role in cancer progression. Our framework assumes that a residue community on a protein structure represents a putative functional subunit of a protein. Thus, high mutation densities in such communities (compared to a random expectation) is very likely to alter protein function. One would expect that mutations influencing residues in these communities will have a high functional impact as they can drive cancer progression. Our observation is consistent with this hypothesis, as we find that missense mutations occupying hotspot communities in proteins structures are highly conserved across species and have a higher molecular functional impact compared to those outside such hotspot communities.

Furthermore, we also observe significantly high enrichment of out putative driver genes with predicted hotspot communities in vital biological processes and pathways that are relevant for oncogenesis. For instance, ontology analysis indicates enrichment of our putative driver genes in biological processes associated with regulation and activation of innate immune response. This observation is consistent with the current notion that dysfunction in immune response contributed through genomic alterations will allow tumor cells to evade immune detection due to lack of effective immune response. Additionally, we also observe a significant enrichment of putative driver genes in cell differentiation and cell growth processes, such as the regulation of hematopoiesis and myeloid cell differentiation, which were previously implicated in tumor growth. Moreover, we observed a high enrichment of our putative driver genes in the regulation of kinase activities including protein serine/threonine and MAP kinase activities. Additionally, these genes are also enriched among ERK1/ERK2 signaling cascade, protein kinase B signaling, PI3K/AKT signaling, FGFR1 signaling, NTRK1 signaling, apoptosis signaling, and various other signaling pathways. Presence of aberrant signaling pathways is an essential hallmark of cancer. Thus, enrichment of our putative genes in critical signaling pathways provides clear biological evidence for their role in cancer. Moreover, these genes are enriched for DNA repair function via non-homologous end joining(NHEJ) and other non-recombination based repair mechanisms. Finally, we note that we observed the same enrichment for the subset of novel genes in our putative driver gene list, that have not been identified as drivers in previous studies.

Genomic alterations that are consequential for tumor growth are often manifested on the transcriptome level such that mutated driver genes are often differentially expressed compared to a healthy population or patients without any mutation in driver genes. We leveraged the transcriptome data from TCGA to further validate out predicted driver genes based on hotspot community identification. We identified 60 genes among our predicted driver genes that were significantly differentially expressed in tumor samples with missense mutations in those genes compared to those without among multiple cancer cohorts. These differentially expressed driver genes include novel as well previously established driver genes. Similar to genetic data, transcriptomic data in TCGA is limited for specific cancer cohort that provides insufficient power to identify all differentially expressed genes. However, we note that 76% of our putative driver genes were differentially expressed in at least one TCGA cancer cohort. These analyses further validate our hotspot community-based driver detection approach.

In the context of investigating the molecular mechanism underlying tumor growth, protein structure-based driver detection methods offer significant advantages over approaches that are only sequence-based. However, structure-based methods suffer from limited coverage of the human proteome. Thus, the applicability of structure-based methods is inherently limited only to mutations that can be mapped onto protein structure. A prior study^36^ has applied homology model derived structures to circumvent the issue of limited structural coverage. However, the accuracy of homology-based models has shown to be limited for various protein complexes and transmembrane proteins. Moreover, modeling protein motions for homology-model derived proteins structures will be most likely less accurate thus affecting the sensitivity of our approach. Nevertheless, significant technical improvement in crystallographic and cryoEM techniques^75^ are expected to expand the current structurally-resolved proteome. In particular, cryoEM technologies^75^ now allows us to obtain a high-resolution structure of large-size proteins and other biomolecular complexes that were previously elusive. Thus, we anticipate an essential role of our approach in future studies aimed at discovering low-frequency drivers in various cancer cohorts. Additionally, knowledge of protein motions (along with structure) can potentially help in uncovering drug interaction with hotspot communities. Such studies are likely to open new therapeutic avenues for various cancers and will help us realize the goal of precision medicine in cancer.

## Acknowledgments

We acknowledge support from the NIH and from the AL Williams Professorship funds. We also acknowledge help of Jonathan Warrell and Timur Galeev for providing valuable feedbacks for improving the manuscript.

