## Supplemental Figures for "Leveraging protein dynamics to identify cancer mutational hotspots in 3D-structures"

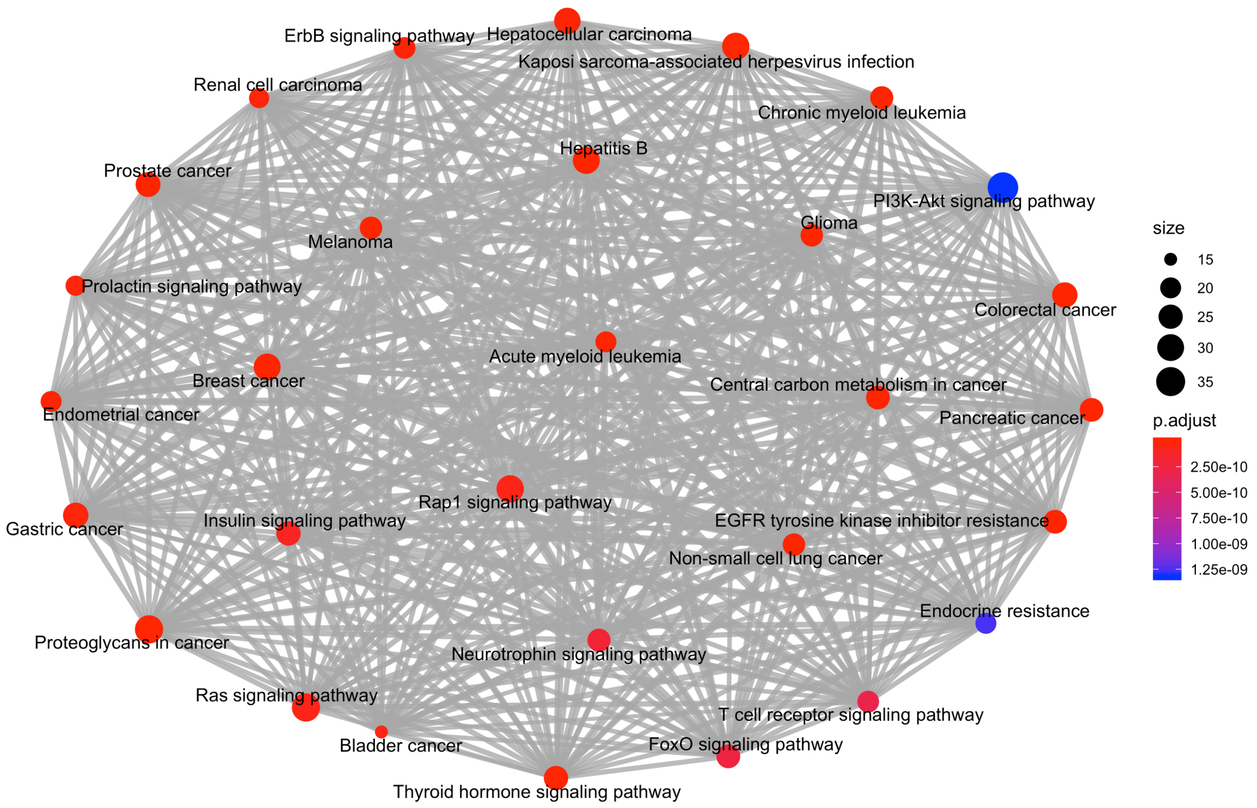


Supplement Fig1: KEGG pathway based enrichment plot for all genes with at least one hotspot community in their corresponding protein three-dimensional structure.

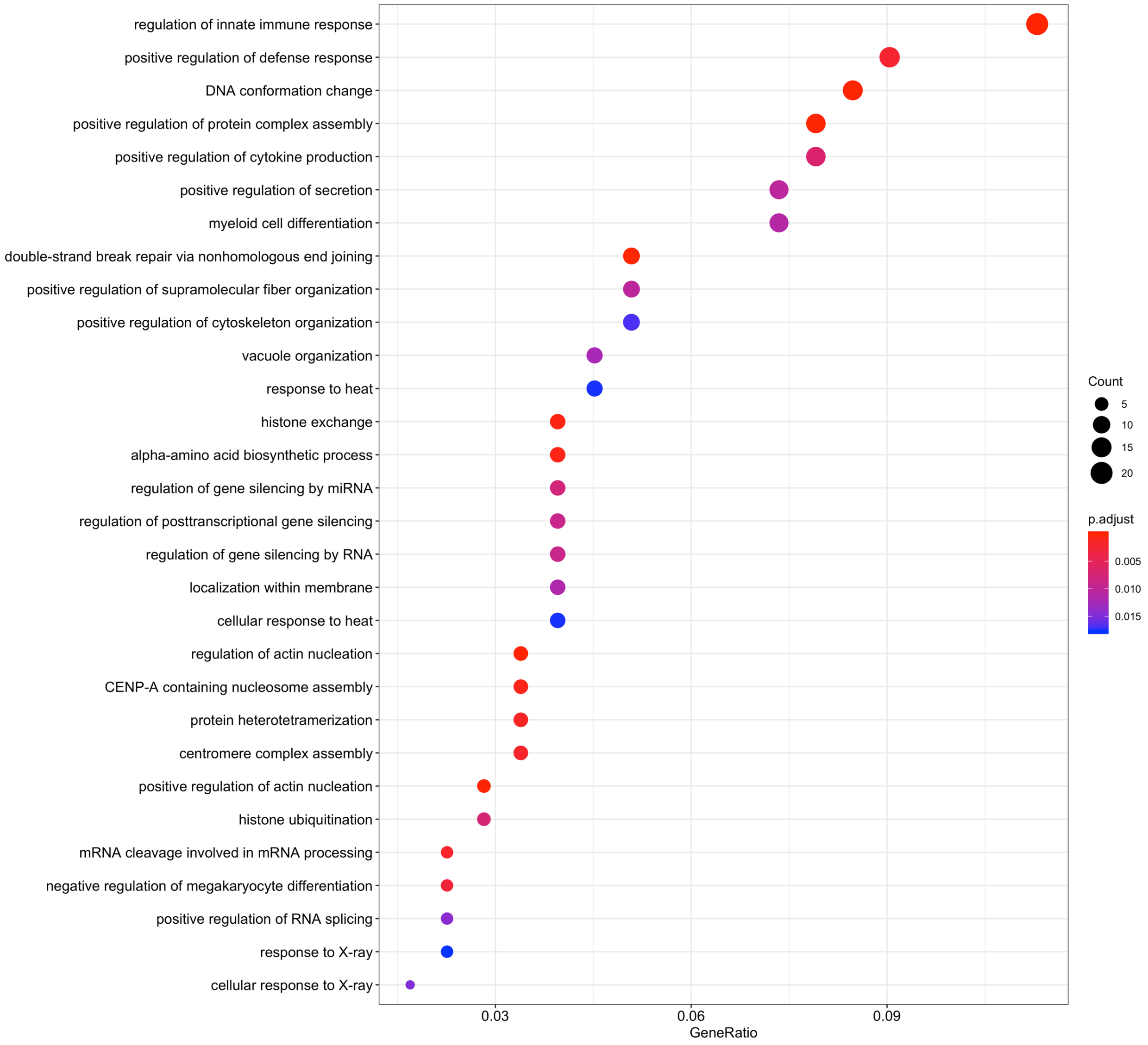


Supplement Fig2: Gene ontology enrichment plot for unique genes (not identified as drivers before) with at least one hotspot community in their corresponding protein three-dimensional structure.
